# Cardiac-sympathetic state predicts action restraint, gated by demonstrated agency

**DOI:** 10.64898/2025.12.29.696920

**Authors:** Neil M. Dundon, Elizabeth J. Rizor, Joanne Stasiak, Jingyi Wang, Viktoriya Babenko, Regina C. Lapate, Scott T. Grafton

## Abstract

Withholding action until the appropriate moment is a core challenge of motivated behavior. Using beat-to-beat cardiac contractility during an incentivized reaching task, we show that cardiac-sympathetic outflow predicts action restraint. Under high-reward conditions that induce a speed–accuracy tradeoff, reduced contractility at the time of instruction preceded premature responses (false starts). Under high-loss-avoidance conditions, elevated pre-movement contractility predicted slower, more controlled initiation, but only among participants with above-median task success. These findings suggest cardiac-sympathetic engagement does not simply serve mobilization but flexibly supports context-appropriate action regulation, with recruitment for restraint gated by demonstrated agency.

## Introduction

Cardiac-sympathetic outflow tracks the demands of effortful engagement. Classic mobilization frameworks demonstrate this principle in tasks requiring fast or active coping, where greater sympathetic drive typically reflects stronger mobilization supporting goal-directed performance (1–3). However, real-world goals are complex, demanding not only energetic mobilization but also precise control over when and whether action is released. A sprinter in the blocks must balance mobilization and *restraint*: maximally primed to explode at the gun, yet suppressing the very movement being prepared; a single premature twitch could mean disqualification.

Converging evidence suggests cardiac-sympathetic outflow can support both sides of this balance. Motor-control circuits can bias toward motivational drive or toward stopping, depending on whether incentives are construed as opportunity or threat. As in the sprinter’s dilemma, this shift can emerge even when the economic stakes and movement demands are held constant (4). Beat-to-beat cardiac studies further suggest that the cardiac-sympathetic system can align with context-dependent control. Across paradigms, sympathetic dynamics covary with behavior and, when neural signals are measured, show context-specific cardiac–cortical coupling (5–7). Notably, these effects are prominent in settings where costs shape policy, including deteriorating environments, decision conflict, and conditions favoring caution over invigoration. Whether cardiac-sympathetic outflow supports mobilization or restraint may depend on perceived control over outcomes. Controllability shapes anticipatory neural processing and subjective threat intensity (8), and agency can govern whether outcomes are credited to one’s own actions and used to guide subsequent action selection and strategy (9, 10).

Here we test whether cardiac-sympathetic outflow predicts both sides of the motivation-restraint balance, and whether its recruitment for restraint depends on the individual’s demonstrated control over outcomes.

## Results

We reanalyzed data from an incentivized reaching task (4), focusing on previously unreported cardiac-sympathetic dynamics indexed by beat-by-beat contractility. The task requires speeded reaches to visual targets under three incentive conditions: high-reward “jackpot”, high-loss-avoidance “robber”, and low-reward “standard” (300 trials; 10% jackpot, 10% robber, 80% standard). The analyzed subset (*n* = 51) showed the same behavioral asymmetry reported previously: jackpot selectively sped movement initiation (cue effect on RT: *F* = 14.7, *p* < .001) while increasing false starts (*F* = 19.9, *p* < .001), consistent with a speed–accuracy tradeoff under high-reward salience. Robber did not speed initiation but instead altered peak acceleration (*F* = 12.6, *p* < .001) and velocity (*F* = 8.4, *p* < .001).

Contractility showed clear event-related structure around the go cue. Contractility significantly increased during the 1 s post-cue interval, followed by an undershoot approximately 2 s post-cue, observed across all conditions (Fig. 1A, right panels). Structure was less evident around the instruction cue (Fig. 1A, left panels).

**Figure 1.**
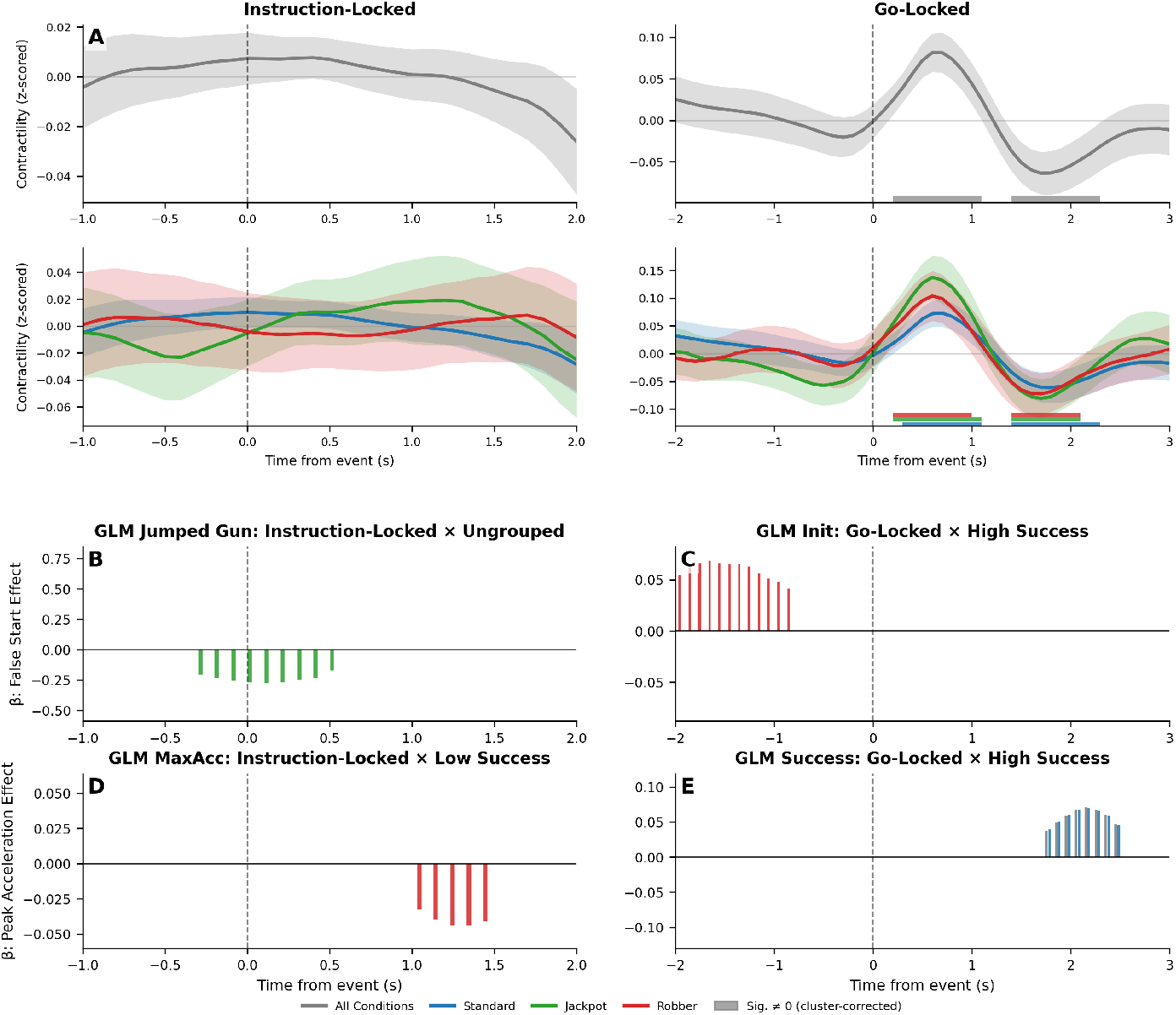
Cardiac contractility dynamics during incentivized reaching. (*A*) Event-locked contractility traces. Top: grand average; bottom: by incentive condition. Left: instruction-locked; right: go cue-locked. Shaded regions: 95% CI; colored bars: significant deviation from zero (cluster-corrected, *p* < 0.05). (*B–E*) Pointwise GLM coefficients (*β* ) for trial-level predictors (cluster-corrected). (*B*) Jackpot trials, all participants: reduced contractility precedes false starts. (*C*) Robber trials, high-success participants: elevated pre-movement contractility predicts slower initiation. (*D*) Robber trials, low-success participants: contractility predicts reduced max acceleration. (*E*) All and standard trials, high-success participants: elevated late contractility on correct trials.

Pointwise GLMs tested whether contractility predicted trial-by-trial behavioral features across the epoch. Two key effects implicated contractility in behavioral restraint. First, on jackpot trials, lower contractility around the instruction cue predicted subsequent false starts (Fig. 1B; cluster *p* < 0.05, −0.3 to 1.5 s relative to cue). This suggests that sympathetic engagement during incentive encoding is associated with successful inhibition of premature responses, particularly under high-reward conditions that induce a behavioral speed–accuracy tradeoff. Second, among participants with above-median success rates, higher contractility preceding the go cue predicted slower movement initiation (Fig. 1C; cluster *p* < 0.05, −1.5 to −0.3 s). This anticipatory cardiac signature of restraint was selective to individuals with superior behavioral control, consistent with perceived controllability shaping cardiac-sympathetic deployment.

Two additional subgroup effects emerged. Among participants with below-median success rates, higher contractility approximately 1 s after the instruction cue predicted reduced peak acceleration (Fig. 1D; cluster *p* < 0.05), suggesting that in individuals with poorer outcome control, elevated sympathetic tone during preparation may dampen rather than support mobilization. Finally, among high-success participants, contractility 2–3 s after the go cue positively predicted trial success on all trial types, and standard trials alone (Fig. 1E; cluster *p* < 0.05). Given its late timing, this effect likely reflects post-movement autonomic correlates of outcome monitoring. No other predictor–condition–subgroup combinations yielded significant clusters at any timepoint.

## Discussion

Classic mobilization frameworks treat cardiac-sympathetic activation as a signature of behavioral mobilization: a coordinated readiness to act, strongest when the organism can actively cope with or control a challenge (1, 2). Our results extend this framework. Sympathetic engagement predicted restraint, not just mobilization, and did so asymmetrically across incentive contexts. Under reward pressure, lower contractility preceded failures to withhold action. Under threat, higher contractility preceded adaptive slowing, but only among individuals whose careful execution had proven effective. This pattern reveals that cardiac-sympathetic outflow is not a unitary mobilization signal but a flexible control resource, associated with context-appropriate action regulation.

Classical models of incentive-motor integration emphasize striatal scaling of movement vigor according to expected value (12, 13). Yet this view underestimates the system’s flexibility: valence, not merely magnitude, determines which circuits dominate motor output (4). The cardiac-sympathetic system offers a complementary window into such regulation. Beat-to-beat contractility provides a continuous readout of central autonomic state (14), and our findings suggest it functions not as generic arousal but as a control signal, recruited when context demands override reward-driven defaults (3, 15). Critically, this recruitment appears gated by demonstrated agency. Individuals who have established control over task outcomes show selective sympathetic engagement under threat, possibly because knowing one *can* succeed makes potential failure feel controllable, amplifying the demand for cautious, well-timed action (9).

Such flexibility is neuroanatomically plausible. Multi-synaptic cortical pathways link motor, cognitive, and affective regions to sympathetic outflow nuclei (16, 17), providing diverse routes through which higher-order systems might recruit cardiac-sympathetic support for distinct control demands. Whether this goal-directed cardiac flexibility mirrors the circuit-level reconfiguration observed in basal ganglia remains to be tested. Concurrent neuroimaging and beat-to-beat cardiac measurement could reveal whether the same cortical controllers that bias striatal output also modulate sympathetic drive, and whether these systems work in concert to support adaptive action regulation.

## Materials and Methods

Task, sample, and behavioral measures are detailed in Dundon et al. (4). Beat-by-beat contractility was recorded using trans-radial electrical bioimpedance velocimetry (TREV) electrodes on the left forearm. We analyzed a subset (*n* = 51) whose TREV waveforms supported stable peak detection after semi-automated preprocessing (11). Contractility was indexed by the peak acceleration of the impedance waveform, a sensitive proxy for sympathetic inotropy. Beat-by-beat estimates were upsampled to 10 Hz, and event-locked epochs were extracted (−1 to 2 s around instruction cue; −2 to 3 s around go cue), each baseline-normalized by subtracting its mean.

Pointwise GLMs employed a two-stage summary statistics approach: ordinary least-squares regression was fit per participant at each timepoint, then aggregated using DerSimonian–Laird random-effects meta-analysis. Three univariate models tested movement initiation time, peak acceleration, and trial success on non-false-start trials; a separate model used binary false-start status across all trials. Inference employed cluster-based permutation testing (10,000 sign-flip permutations) with cluster-forming threshold |*z*| > 1.96, controlling family-wise error at cluster level.

## Data Availability

Anonymized data underlying this study will be deposited in a community-recognized, persistent repository and made publicly available no later than the time of publication of the final peer-reviewed article. Data, code, protocols, and key materials used and generated in this study are unambiguously identified and will be made available in accordance with ASAP’s Open Science Policy, which requires that research outputs be deposited in community-recognized repositories with information to facilitate reuse and licenses that allow reuse. All data and materials will be assigned persistent identifiers (e.g., DOI) to facilitate linking and citation.

## Acknowledgments

The authors thank Mario Mendoza at UCSB’s Brain Imaging Center for data collection assistance, and Taylor Li, Kiana Sabugo, Christina Villanueva, Parker Barandon, Renee Beverly-Aylwin, and Alexandra Stump for contributions to data collection and processing. This research was funded by Aligning Science Across Parkinson’s ASAP-020-519 through the Michael J. Fox Foundation for Parkinson’s Research (MJFF). For the purpose of open access, the authors have applied a CC BY public copyright license to all Author Accepted Manuscripts arising from this submission.

